# Targeting the Mitochondrial ACSS1-Dependent Acetate–Pyrimidine Axis Suppresses Mantle Cell Lymphoma Progression

**DOI:** 10.1101/2025.08.29.673065

**Authors:** Johnvesly Basappa, Aaron R. Goldman, Cosimo Lobello, Shengchun. Wang, David Rushmore, Olga Melnikov, Neil V. Sen, Vinay S. Mallikarjuna, Priyanka Jain, Masoud Edalati, David S Nelson, Kathy Q. Cai, Pin Lu, Reza Nejati, Hossein Borghaei, Pradeep K Gupta, Kavindra Nath, Kathryn E. Wellen, Mariusz A. Wasik

## Abstract

Acetate serves as an alternative carbon source in nutrient-limited tumors, yet its role in supporting nucleotide biosynthesis remains poorly understood. Here, we identify the mitochondrial enzyme ACSS1 as a key metabolic driver in mantle cell lymphoma (MCL), diffuse large B-cell lymphoma (DLBCL), and chronic lymphocytic leukemia (CLL). ACSS1 is frequently overexpressed and catalyzes the conversion of acetate to mitochondrial acetyl-CoA, sustaining oxidative metabolism and biosynthesis under nutrient stress. Genetic silencing of ACSS1 impairs mitochondrial respiration and disrupts acetate incorporation into acetyl-CoA, TCA cycle intermediates, glutamate, and aspartate, while markedly reducing ^13^C-acetate labeling of dihydroorotate and orotate, intermediates in de novo pyrimidine synthesis. Untargeted metabolomics reveal enrichment of pyrimidine biosynthesis pathways in ACSS1-high cells. Notably, acetate or uridine supplementation rescues the growth of ACSS1-deficient cells, confirming a functional link between acetate metabolism and nucleotide synthesis. Importantly, *in vivo* studies using luciferase-labeled JeKo-1 and Maver mantle cell lymphoma xenografts demonstrate that ACSS1 knockdown significantly suppresses tumor growth. NSG mice injected with ACSS1-silenced cells exhibit a marked reduction in tumor burden, as measured by bioluminescence imaging and total photon flux, with significant differences observed at days 14 and 21 post-injection. These findings establish that ACSS1 is required not only for metabolic adaptation *in vitro* but also for lymphoma progression *in vivo*. Collectively, our results uncover an ACSS1-dependent mitochondrial acetate–pyrimidine axis that sustains lymphoma growth and represents a previously unrecognized therapeutic vulnerability.

**Statement of Significance:** This study identifies ACSS1 as a critical metabolic vulnerability in mantle cell lymphoma (MCL), linking mitochondrial acetate metabolism to de novo pyrimidine biosynthesis and tumor progression. We demonstrate that ACSS1 is frequently overexpressed in MCL and is essential for converting acetate into mitochondrial acetyl-CoA, thereby sustaining TCA cycle activity, nucleotide production, and cell survival under nutrient stress. Loss of ACSS1 disrupts this acetate–pyrimidine axis, impairing oxidative metabolism and reducing lymphoma cell viability in vitro. Importantly, ACSS1 silencing significantly suppresses tumor growth in vivo, establishing its requirement for lymphoma progression. The ability of acetate or uridine supplementation to rescue ACSS1-deficient cells further highlights the functional coupling between mitochondrial acetate utilization and nucleotide synthesis. Together, these findings reveal a previously unrecognized mechanism of metabolic adaptation in aggressive lymphomas and offer ACSS1-mediated acetate metabolism as a promising therapeutic target.

## Introduction

In higher vertebrates, the swift growth of cells mainly relies on the intake of glucose and glutamine(1). Acetate is another nutritional source vital for the progression of many cancers, which often exhibit dysregulated metabolism, and it is essential under nutrient-deprived conditions(2). The mitochondrial acetyl-CoA synthetase short-chain family member 1 (ACSS1) catalyzes acetate, an energy source during nutrient deprivation conditions, to produce mitochondrial acetyl-CoA(2, 3). A recent study suggests that acetate promotes PD-L1 expression and immune evasion by upregulating c-Myc(4). Indeed, in ketogenic conditions, such as prolonged fasting or diabetes, the liver also generates sufficient quantities of acetate and releases it into the bloodstream(5). Mammals have three isoforms of the short-chain acetyl-CoA synthetases family: ACSS1, ACSS2, and ACSS3, which convert acetate into acetyl-CoA(6, 7), which is essential for cell growth and proliferation(8, 9). ACSS1 and ACSS3 localize in the mitochondria, while ACSS2 is found in the nucleus and cytoplasm(10). Among the three isoforms, ACSS2’s role in acetate metabolism and cancer progression has been extensively studied in various types of cancer, including pancreatic cancer(11), hepatocellular carcinoma(12, 13), glioblastoma(14), breast cancer(15), prostate cancer(15), bladder cancer(16) and cervical squamous cell carcinoma(17). ACSS1 plays a crucial role in thermogenesis during fasting, in low-glucose or ketogenic states, and is vital for survival(18). The expression of ACSS1 is regulated in an estrogen receptor-dependent manner, and the estrogen hormone plays a role in regulating the incorporation of acetate into DNA(19, 20). In highly oxygen-rich environments, most acetyl-CoA is generated via the glucose pathway. Conversely, during low oxygen conditions, acetate serves as a source for acetyl-CoA production(21). The exact function of mitochondrial ACSS1 in acetate metabolism within lymphoma remains unknown.

Mantle cell lymphoma (MCL) is a type of B-cell lymphoma, characterized by a median overall survival of just 3-5 years (22). While a subset of patients with mantle cell lymphoma (MCL) initially respond to Bruton tyrosine kinase (BTK) inhibitors, resistance, either intrinsic or acquired, is a common clinical challenge. Emerging evidence suggests that metabolic reprogramming toward oxidative phosphorylation (OXPHOS) contributes to disease progression and therapeutic resistance in MCL. In this study, we employed MCL cell lines as a model to investigate the role of ACSS1 in acetate metabolism under nutrient-deprived conditions. Using extensive ^13^C-acetate stable isotope tracing and genetic knockdown of ACSS1, we uncover a previously unrecognized function of ACSS1 in supporting acetate-fueled survival and de novo pyrimidine synthesis. These findings highlight ACSS1 as a critical mediator of metabolic adaptation in aggressive, therapy-resistant MCL.

## Results

### ACSS1 expression shows variable expression in MCL and DLBCL samples

To validate ACSS1 protein expression in clinical samples, we performed immunohistochemistry (IHC) on tumor tissue microarrays (TMAs) from mantle cell lymphoma (MCL) patient biopsies (n = 27) obtained from Fox Chase Cancer Center. ACSS1 staining was detected in 92.6% of MCL cases (25/27), with expression levels categorized as strong in 7 cases (25.9%), moderate in 9 cases (33.3%), and weak in 9 cases (33.3%) (Fig. 1A, B). A representative low-magnification image of the TMA block is shown in Supplementary Fig. 1A. In chronic lymphocytic leukemia (CLL) patient samples (n = 13), IHC analysis revealed ACSS1 expression in 7 cases (53.8%), as shown in Supplementary Fig. 1A. We next evaluated diffuse large B-cell lymphoma (DLBCL) samples (n = 44). We observed ACSS1 positivity in 35 cases (79.5%) (Fig. 1C), including strong expression in 6 cases (13.6%), moderate in 14 (31.8%), and weak in 15 (34.1%) (Fig. 1D). The corresponding TMA overview is provided in Supplementary Fig. 1B. These data confirm frequent ACSS1 protein expression across multiple B-cell malignancies, consistent with transcriptomic analyses.

**Fig. 1.**
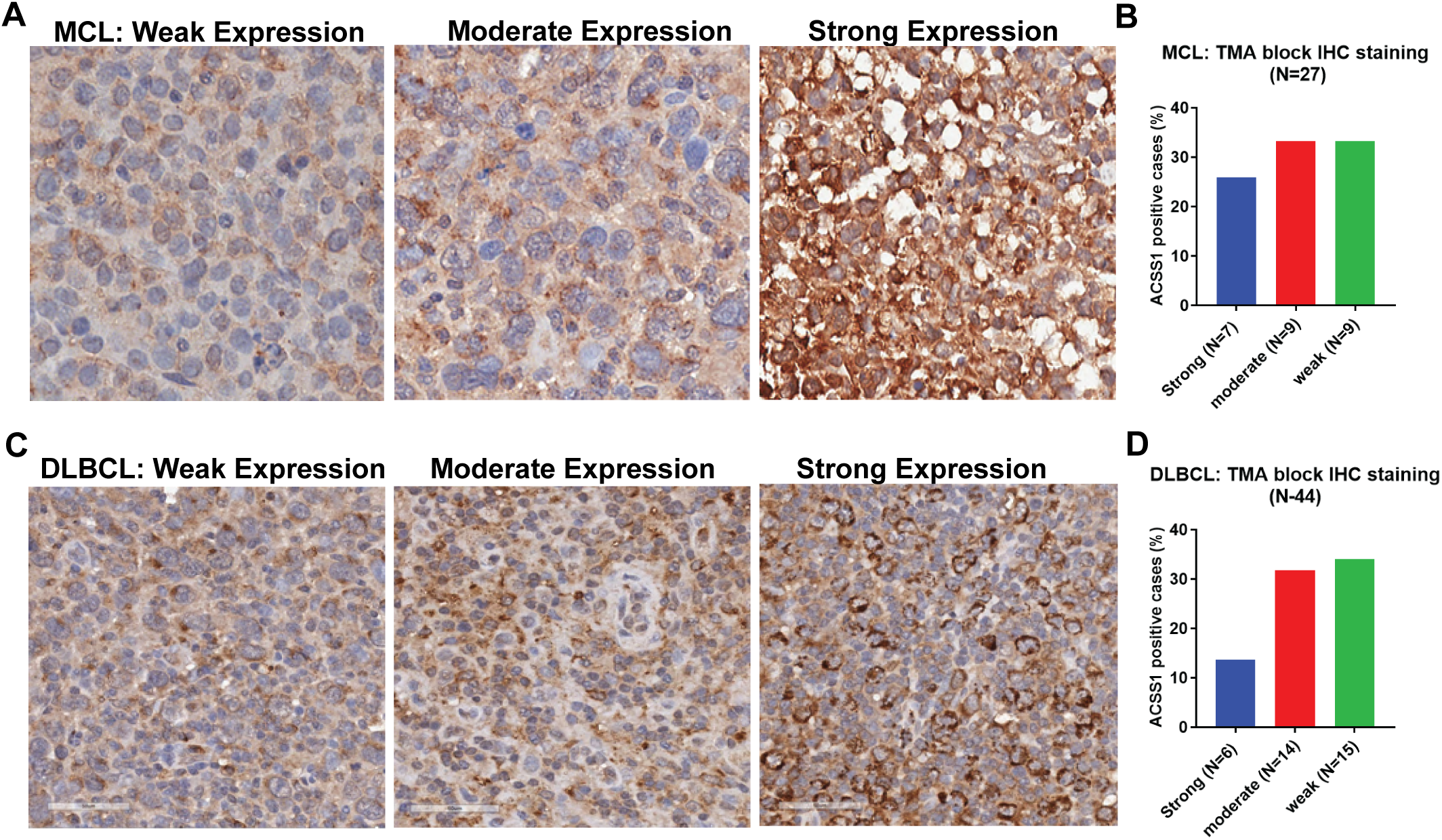
ACSS1 expression shows variable expressions in MCL and DLBCL samples. **A,** Representative immunohistochemistry (IHC) images of ACSS1 protein expression in mantle cell lymphoma (MCL) tissue microarray (TMA) cores, showing examples of strong, moderate, and weak staining. **B**, Quantification of ACSS1 IHC staining intensity in MCL (n = 27) samples. Staining intensity was categorized as strong, moderate, weak, or negative. **C,** Representative IHC images of ACSS1 expression in DLBCL TMA cores, corresponding to varying expression levels. **D,** Quantification of ACSS1 IHC staining intensity in diffuse large B-cell lymphoma (DLBCL, n = 44) samples. Staining intensity was categorized as strong, moderate, weak, or negative.

To validate the specificity of the ACSS1 antibody for IHC staining, a breast cancer sample was tested using immunofluorescence with ACSS1 and the mitochondrial marker VDAC1(Supplementary Fig. 2A). The negative control shows no signal for ACSS1 staining (Supplementary Fig. 2B). To confirm ACSS1 expression in mitochondria, we performed fractionation on MCL cell lines Maver and Jeko-1. Immunoblotting revealed that ACSS1 is primarily located in mitochondria (Supplementary Fig. 2C).

To further investigate the ACSS1 expression in various cancer models, we analyzed publicly available RNA-seq data from hematological malignancy cell lines using the Expression Atlas (https://www.ebi.ac.uk/gxa/home). This analysis focused on hematopoietic and lymphoid-derived cancer models, including mantle cell lymphoma (MCL), diffuse large B-cell lymphoma (DLBCL), chronic lymphocytic leukemia (CLL), acute myeloid leukemia (AML), and anaplastic large cell lymphoma (ALCL). Among acetyl-CoA synthetases, mitochondrial ACSS1 exhibited notably high expression in non-Hodgkin lymphoma (NHL) cell lines and patient-derived samples. In contrast, ACSS3 expression was negligible across all profiled lines (Supplementary Fig. 3A, B). To extend these observations, we examined the expressions of ACSS1 and ACSS2 in hematopoietic and lymphoid malignancies using The Cancer Genome Atlas (TCGA). ACSS1 was overexpressed in 36.2% of 221 hematopoietic and lymphoid samples, with only 1.36% showing reduced expression (Table 1). Given the limited characterization of ACSS1 in solid tumors, we further assessed its expression across multiple cancer types using TCGA. ACSS1 was significantly overexpressed relative to matched normal tissue in several malignancies, including lung (10.89%), ovarian (29.32%), prostate (16.47%), central nervous system (79.91%), cervical (84.04%), large intestine (25.9%), and head and neck squamous cell carcinoma (24.71%) (Table 1). These findings suggest that ACSS1 upregulation is a recurrent feature across both hematological and solid tumors, highlighting its potential role in cancer-associated acetate metabolism.

**Table 1.** ACSS1 expression profile in the TCGA database across cancer models.

| Cancer type | Total number of samples tested | No. of samples over expressed | Over expressed (%) | No. of samples under expressed | Under expressed (%) |
| --- | --- | --- | --- | --- | --- |
| Hematologic and lymphoid | 221 | 80 | 36.2 | 3 | 1.36 |
| Central Nervous System | 697 | 557 | 79.91 | 32 | 4.59 |
| Lung | 1019 | 111 | 10.89 | None | None |
| Ovary | 266 | 78 | 29.32 | None | None |
| Cervix | 307 | 258 | 84.04 | 29 | 9.45 |
| Large Intestine | 610 | 158 | 25.9 | 12 | 1.97 |
| Prostate | 498 | 82 | 16.47 | 3 | 0.6 |
| Head and Neck | 522 | 129 | 24.71 | None | None |

### Acetate feeds the TCA cycle in MCL and DLBCL cells in an ACSS1-dependent manner

Our analysis of patient-derived TMA, TCGA data, and hematological cancer cell lines revealed variable ACSS1 expression across mantle cell lymphoma (MCL) and diffuse large B-cell lymphoma (DLBCL) models. Notably, ACSS1 mRNA levels were low in MCL-RL cells but were significantly elevated in Jeko-1 and Maver MCL cell lines, exhibiting 18-fold and 24-fold higher expression, respectively (Fig. 2A). In contrast, ACSS2 expression remained low and was not differentially expressed across these MCL lines (Fig. 2B). Similarly, high ACSS1 mRNA levels were observed in DLBCL cell lines, whereas one cell line OCI-LY1 displayed low ACSS1 expression (Fig. 2C). Consistent with MCL, ACSS2 expression was uniformly low in DLBCL lines (Fig. 2D). Immunoblot analysis of five MCL cell lines (Jeko-1, Maver, Mino, Granta-519, and RL) confirmed robust ACSS1 protein expression in all except MCL-RL cells (Fig. 2E). Likewise, immunoblotting of four DLBCL cell lines demonstrated elevated ACSS1 protein levels in all except OCI-LY1 cells (Fig. 2F). These findings suggest that ACSS1 expression correlates with variable expressions in both MCL and DLBCL models.

**Fig. 2.**
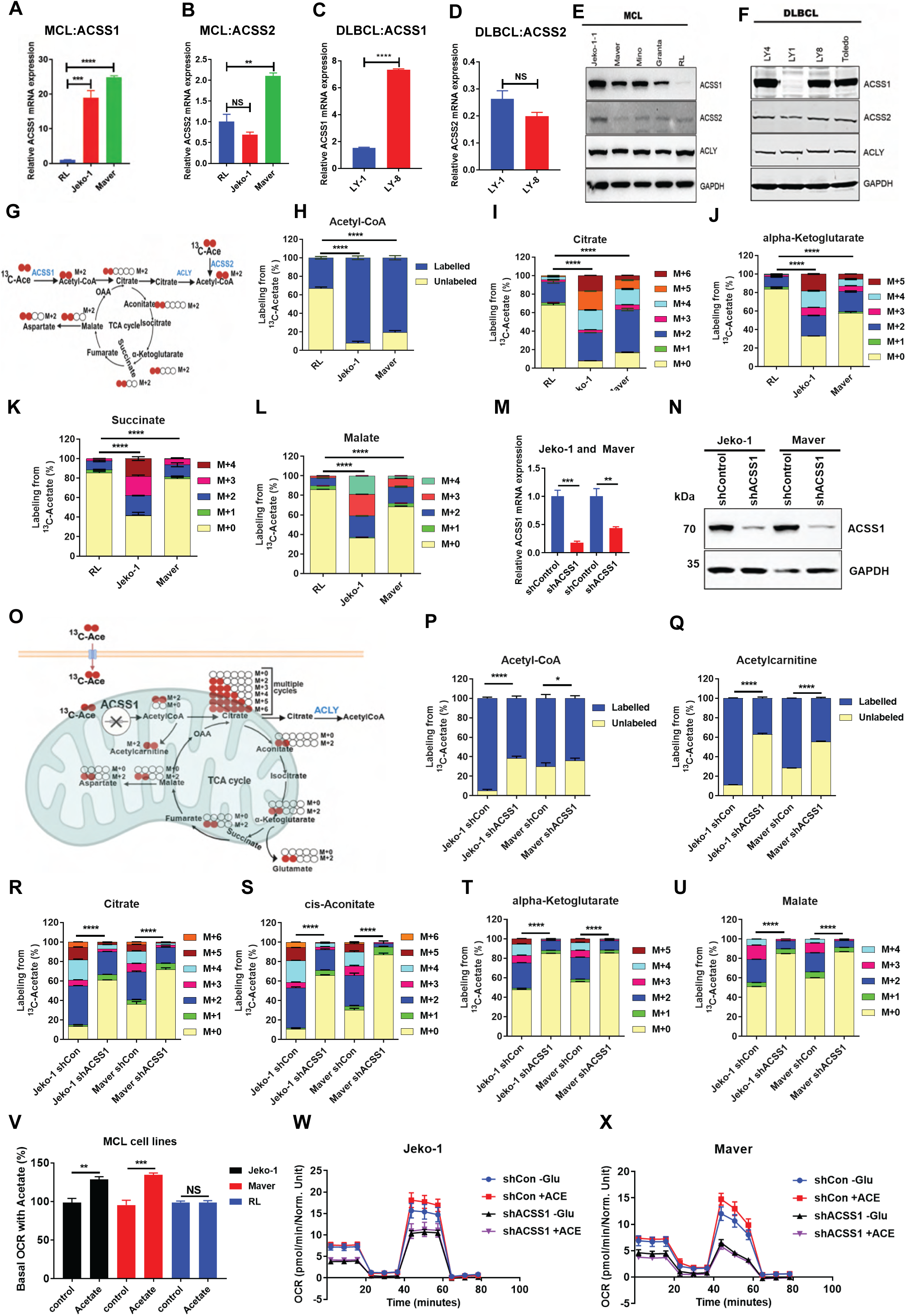
Acetate feeds the TCA cycle in MCL and DLBCL cells in an ACSS1-dependent manner. **A-D**, Quantitative RT-PCR analysis of ACSS1 and ACSS2 mRNA expression in MCL (RL, Jeko-1, Maver) and DLBCL (OCI-LY1, OCI-LY8) cell lines. ACSS1 is highly expressed in Jeko-1, Maver, and OCI-LY8, while ACSS2 remains uniformly low. **E, F**, Immunoblot analysis of ACSS1 protein levels across five MCL (e) and four DLBCL (f) cell lines confirms elevated expression in all except RL and OCI-LY1, respectively. **G,** A schematic of ¹³C-acetate isotope tracing. **H, I,** ¹³C-acetate isotope tracing and LC-MS analysis of TCA cycle intermediates in RL, Jeko-1, and Maver cells. unalabeled and labeled acetyl-CoA (**H**), M+0 to M+n isotopologues of citrate (**I**), α-ketoglutarate (**J**), succinate (**K**), and malate (**L**) are enriched in ACSS1-high lines. **M, N**, ACSS1 knockdown efficiency confirmed by qPCR and immunoblotting in Jeko-1 and Maver cells. **O**, Schematic of isotope tracing strategy using ¹³C-acetate in glucose- and glutamine-deprived media. **P, Q**, ACSS1 knockdown reduces ^13^C-acetate derived labeling of acetyl-CoA (**P**) and mitochondrial acetylcarnitine (**Q**), confirming ACSS1’s role in mitochondrial acetate conversion. **R–U**, Stable isotope tracing reveals significantly decreased ¹³C incorporation into citrate (**R**), aconitate (**S**), α-ketoglutarate (**T**), and malate (**U**) upon ACSS1 silencing. **V**, Basal oxygen consumption rate (OCR) increases in ACSS1-high Jeko-1 and Maver cells after acetate supplementation; no effect in low-ACSS1 RL cells. **W, X**, ACSS1 knockdown reduces basal and stress-induced OCR in Jeko-1 (u) and Maver (v) cells under acetate-reliant conditions. All graphs show mean ± SD (n = 3 biological replicates), and all statistical analyses were conducted with unpaired t-tests: ^∗^p < 0.05, ^∗∗^p < 0.01, ^∗∗∗^p < 0.001, ^∗∗∗∗^p < 0.0001

To investigate the relationship between ACSS1 expression and acetate metabolism, we performed stable isotope tracing using ^13^C-labeled acetate, followed by liquid chromatography–mass spectrometry (LC-MS) to analyze mass isotopologue distributions (MIDs) of tricarboxylic acid (TCA) cycle intermediates as illustrated in schematics (Fig. 2G). Stable isotope tracing with ^13^C-labeled acetate revealed distinct differences in acetate utilization correlated with ACSS1 expression levels. Analysis of acetyl-CoA mass isotopologue distributions showed significant variation in both unlabeled and labeled species among the three MCL cell lines (Fig. 2H). To further assess the metabolic fate of acetate-derived carbon, we quantified the incorporation of ^13^C into tricarboxylic acid (TCA) cycle intermediates, including citrate, α-ketoglutarate (α-KG), succinate, and malate. Citrate labeling patterns differed markedly among RL, Jeko-1, and Maver cells, with Jeko-1 and Maver cells exhibiting significantly increased ^13^C-acetate–derived labeling compared with RL cells (p < 0.0001) (Fig. 2I). Similarly, α-KG showed significant differences in M+2, M+3, M+4, and M+5 isotopologues, particularly between Jeko-1/Maver and the low-ACSS1 RL cells (Fig. 2J). These trends were also observed in succinate (Fig. 2K) and malate (Fig. 2L), with higher ^13^C incorporation in Jeko-1 and Maver relative to RL. Collectively, these findings indicate that elevated ACSS1 expression enhances acetate utilization and contributes to increased TCA cycle flux in MCL cell lines.

To elucidate the role of ACSS1 in mitochondrial acetate metabolism, we silenced ACSS1 in two MCL cell lines, Jeko-1 and Maver, which exhibit high endogenous ACSS1 expression. Knockdown efficiency was confirmed at the transcript and protein levels using quantitative PCR and immunoblotting, respectively (Fig. 2M, N). To determine whether ACSS1 drives mitochondrial acetate metabolism, we conducted ^13^C-acetate labeling analysis in both shControl and ACSS1 knockdown (shACSS1) cell lines, as illustrated in the schematic (Fig. 2O). Given that most mammalian cells rely primarily on glucose and glutamine for energy production, we hypothesized that ACSS1 is particularly critical under nutrient-deprived conditions. Cells were cultured for 24 hours in ^13^C-acetate–supplemented RPMI medium lacking glucose and glutamine, with 1% dialyzed FBS. Metabolomic analysis revealed a significant reduction in acetyl-CoA labeling in ACSS1 knockdown Jeko-1 cells p< 0.0001(Fig. 2P). To distinguish between cytosolic and mitochondrial acetyl-CoA pools, we quantified mitochondrial acetylcarnitine labeling. ACSS1 knockdown led to a marked reduction in acetylcarnitine M+2 labeling in both Jeko-1 (>40%; p < 0.0003) and Maver ( p < 0.0005) cells (Fig. 2Q), supporting a key role for ACSS1 in mitochondrial acetate utilization. Further analysis of ^13^C-acetate–derived MIDs in TCA cycle intermediates demonstrated that ACSS1 knockdown decreased citrate M+2–M+5 labeling in both cell lines, consistent with reduced carbon entry from acetate through multiple TCA cycle turns (Fig. 2R). Additional intermediates including aconitate, α-ketoglutarate, and malate exhibited similar reductions in ^13^C labeling upon ACSS1 silencing (Fig. 2S-U). These results collectively indicate that ACSS1 is essential for converting acetate into mitochondrial acetyl-CoA, fueling oxidative phosphorylation via the TCA cycle, particularly under nutrient-limited conditions. Thus, ACSS1 supports mitochondrial bioenergetics in MCL cells with elevated ACSS1 expression.

Building on the observed role of ACSS1 in mitochondrial acetate metabolism and its influence on TCA cycle activity, we hypothesized that ACSS1 also modulates oxidative phosphorylation (OXPHOS) through regulation of oxygen consumption rate (OCR). To evaluate this, we measured basal OCR in glucose-starved RL (low ACSS1-expressing), Jeko-1, and Maver (high ACSS1-expressing) MCL cell lines following exposure to 1 mM acetate. Acetate supplementation led to a significant increase in OCR in Jeko-1 (24%, *P* < 0.002) and Maver (30%, *P* < 0.0001) cells. In contrast, RL cells exhibited minimal response (Fig. 2V). These findings suggest that ACSS1 expression enables acetate-fueled mitochondrial respiration under glucose-limiting conditions. To further confirm the role of ACSS1 in regulating acetate-dependent OXPHOS, we compared OCR in control (shCon) and ACSS1 knockdown (shACSS1) Jeko-1 and Maver cells. In both cell lines, ACSS1 silencing resulted in a marked reduction in basal and stress-induced OCR compared to control cells (Fig. 2W, X), indicating that ACSS1 is critical for sustaining mitochondrial respiration in the presence of acetate. Collectively, these results underscore the importance of ACSS1 in driving acetate-supported oxidative metabolism in MCL cells.

### ACSS1 supports pyrimidine metabolism

Based on the differential expression of ACSS1 in MCL cell lines high in Jeko-1 and Maver, and low in RL we aimed to investigate ACSS1’s role in acetate metabolism under nutrient-deprived conditions. To do this, we performed ^13^C-acetate tracing in these cell lines. Our metabolic analysis, which examined changes in steady-state metabolite pathways and enrichment patterns, revealed perturbations in pyrimidine biosynthesis pathways. To determine whether acetate directly contributes to pyrimidine biosynthesis, we analyzed the incorporation of ^13^C into key metabolic precursors, as illustrated in the schematic (Fig. 3A). In Jeko-1 and Maver cells, which express high levels of ACSS1, we observed a significant increase in ^13^C-acetate–derived M+2 to M+5 labeling of aspartate and glutamate two critical intermediates for de novo pyrimidine synthesis compared to low-ACSS1-expressing RL cells (P < 0.0001, Fig. 3B, C). These findings indicate that ACSS1 enhances pyrimidine biosynthesis by supporting the generation of TCA cycle–derived intermediates under acetate-reliant, nutrient-limited conditions. To further evaluate acetate’s contribution to pyrimidine synthesis, we examined the labeling patterns in downstream intermediates. Isotope tracing revealed a significant enrichment of M+2 to M+5 ^13^C-labeled orotate, UMP, UDP, and CDP in Jeko-1 and Maver cells compared to RL (P < 0.0001, Fig. 3D–G), further supporting a model in which ACSS1 drives acetate-dependent pyrimidine biosynthesis.

**Fig. 3.**
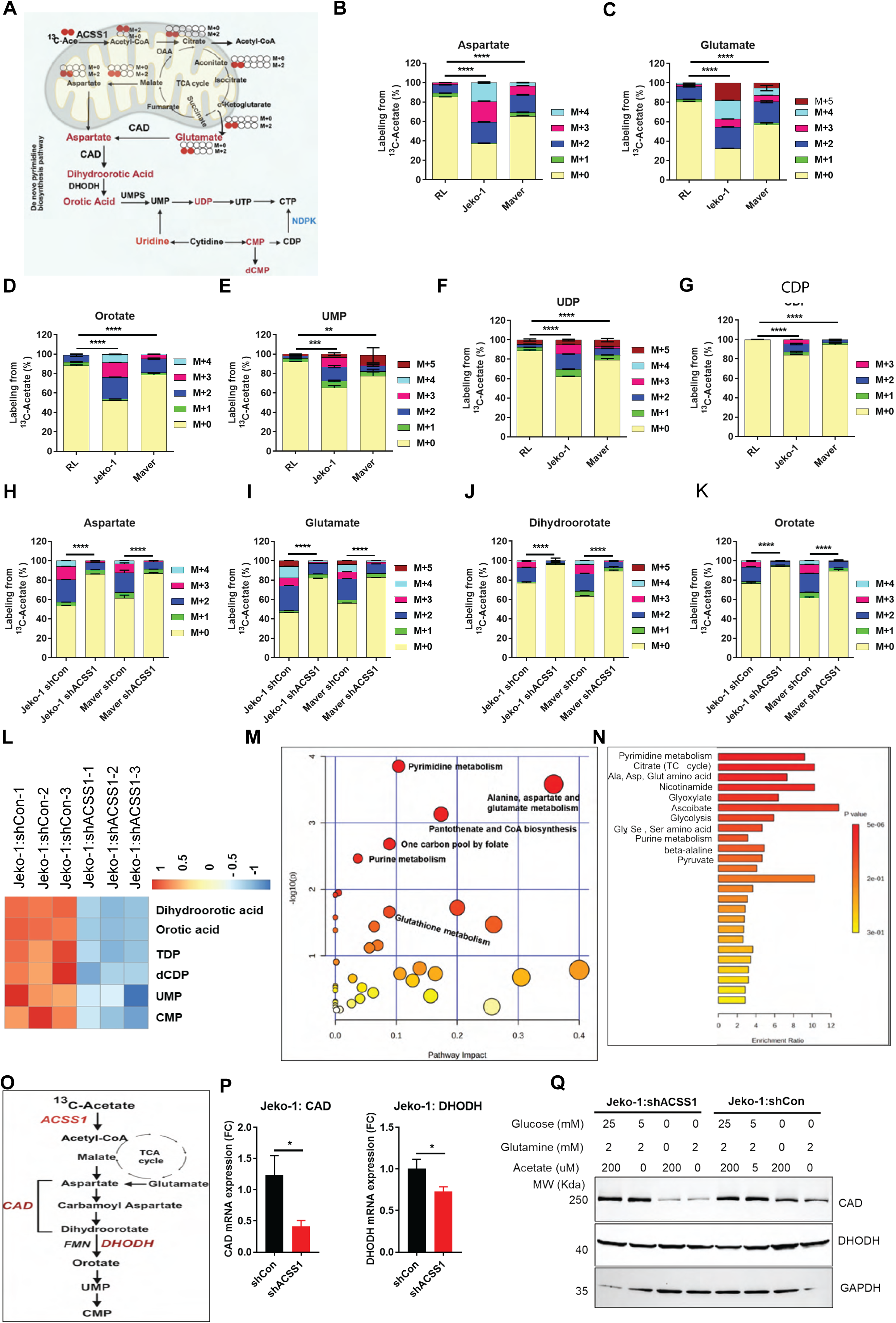
ACSS1 supports pyrimidine metabolism. **A**, Schematic illustrating ¹³C-acetate tracing strategy to assess incorporation into TCA cycle intermediates and downstream pyrimidine biosynthesis. **B, C**, Mass isotopologue distribution analysis of aspartate (**B**) and glutamate (**C**) in RL (low ACSS1), Jeko-1, and Maver (high ACSS1) MCL cell lines shows significantly increased ¹³C-labeling in ACSS1-high cells under nutrient-limited conditions (RPMI lacking glucose/glutamine, supplemented with ¹³C-acetate). **D–G**, Enhanced incorporation of ¹³C into pyrimidine intermediates orotate (**D**), UMP (**E**), UDP (**F)**, and CDP (**G**) in Jeko-1 and Maver compared to RL cells, indicating acetate contribution to de novo pyrimidine synthesis. **H, I**, Knockdown of ACSS1 via shRNA (shACSS1) in Jeko-1 and Maver cells reduces ¹³C-labeling of aspartate (**H**) and glutamate (**I**) compared to scramble controls (shCon). **J, K**, ¹³C-labeling of dihydroorotate (**J**) and orotate (**K**) is also significantly decreased in shACSS1 cells, confirming impaired pyrimidine biosynthesis. **L**, Heat map highlighting significantly altered pyrimidine-related metabolites in Jeko-1 shACSS1 vs. shCon cells (fold change >1.5, P < 0.05). **M, N**, Metabolite enrichment and pathway analysis of 206 profiled metabolites in shACSS1 cells reveals prominent suppression of the pyrimidine synthesis pathway. **O**, Schematic of pyrimidine biosynthesis pathway showing ACSS1’s proposed role and key enzymes (CAD, DHODH). **P**, qRT-PCR analysis of CAD and DHODH mRNA levels shows reduced expression in shACSS1 cells. **Q**, Western blot analysis reveals decreased CAD protein expression in shACSS1 cells cultured in acetate/glutamine-only media, supporting a regulatory role of ACSS1 in pyrimidine biosynthetic enzyme expression under metabolic stress. All graphs show mean ± SD (n = 3 biological replicates), and all statistical analyses were conducted with unpaired t-tests: ^∗^p < 0.05, ^∗∗^p < 0.01, ^∗∗∗^p < 0.001, ^∗∗∗∗^p < 0.0001.

Next, we hypothesized that overexpression of ACSS1 in Jeko-1 and Maver cell lines drives acetate metabolism in MCL. To investigate this, we silenced ACSS1 expression in both cell lines using lentiviral short hairpin RNA (shRNA) targeting ACSS1. The resulting shACSS1 and shCon cell lines were subsequently used in ^13^C-acetate tracing experiments to assess alterations in acetate metabolism. Our metabolomics analysis data showed a highly significant reduction in M+2 to M+5 ^13^C-labeling of aspartate and glutamate in shACSS1 Jeko-1 and Maver cells compared to shCon (P < 0.0001, Fig. 3H, I). Next, ^13^C-acetate isotope tracing revealed a significant reduction in M+2 to M+5 ^13^C-labeled dihydroorotate and orotate in shACSS1 Jeko-1 and Maver cells compared to scramble controls (P < 0.0001, Fig. 3J, K). Comparative analysis of shACSS1 and shCon Jeko-1 and Maver cells cultured with ^13^C-acetate revealed significant alterations in cellular metabolism. Of the 206 metabolites analyzed, 46 metabolites were significantly changed (absolute fold change >1.5, P < 0.05), with the majority associated with the pyrimidine synthesis pathway (Supplementary Fig. 4A, B). A subset of altered pyrimidine metabolites from Jeko-1 cells is highlighted in Fig. 3L. Additionally, pathway and enrichment analyses confirmed the enrichment of metabolites linked to pyrimidine biosynthesis (Fig. 3M, N). These findings suggest that ACSS1 plays a critical role in channeling acetate-derived carbon into TCA cycle intermediates that support de novo pyrimidine synthesis.

We hypothesized that silencing ACSS1 in Jeko-1 cells would impact the expression of key enzymes involved in pyrimidine biosynthesis, as illustrated in the schematic (Fig. 3O). To test this, we isolated RNA from shCon and shACSS1 cells cultured in complete media and performed quantitative RT-PCR to assess the expression of CAD and DHODH. The results demonstrated altered expression of both genes in shACSS1 cells compared to shCon (Fig. 3P). To further evaluate these findings at the protein level, we conducted Western blot analysis under different nutrient conditions. Notably, shACSS1 cells exhibited reduced CAD protein expression when cultured in acetate/glutamine-only media, whereas shCon cells maintained stable CAD levels (Fig. 3Q).

### ACSS1 supports growth under nutrient restriction

We confirmed efficient ACSS1 knockdown (KD) in Maver and Jeko-1 cells using Western blot analysis, with two of four shRNA clones achieving near-complete protein depletion (Fig. 4A). ACSS1 KD led to a significant reduction in cell growth (P < 0.0001), as assessed by automated cell counting and trypan blue exclusion (Fig. 4B). To assess the role of ACSS2, Jeko-1 and Maver cells were treated with an ACSS2 inhibitor (10–30 µM), resulting in a dose-dependent decrease in cell growth (Fig. 4C). Under metabolic stress conditions, cells were cultured in glucose- and glutamine-free media supplemented with 1% dialyzed FBS. While the viability of shCon and shACSS1 cells remained stable in complete media, ACSS1 KD cells showed a marked decline in viability under nutrient deprivation. Supplementation with 100 µM acetate significantly rescued viability in ACSS1 KD cells (from 16% to >68%) and in controls (from 36% to >82%), highlighting ACSS1’s critical role in acetate metabolism (Fig. 4D, E). Although glucose and glutamine supplementation restored viability in control cells, glutamine was less effective in rescuing ACSS1 KD cells. To determine whether impaired pyrimidine synthesis contributed to cell death, we supplemented cultures with 100 µM uridine under nutrient-deprived conditions. Uridine markedly enhanced viability in both control and ACSS1 KD cells (P < 0.0001; Fig. 4F, G). A schematic model (Fig. 4H) illustrates how ACSS1 overexpression promotes the generation of TCA cycle intermediates from mitochondrial acetate, providing a critical carbon source that enhances nucleotide biosynthesis and supports tumor growth. In contrast, ACSS1 knockdown cells (Fig. 5I) exhibit impaired production of mitochondrial acetate-derived TCA cycle intermediates in the absence of acetate, leading to increased cell death and reduced tumor burden. Notably, acetate supplementation rescues cell death in ACSS1-deficient cells. This compensatory effect is mediated by upregulation of the cytosolic acetate-metabolizing enzyme ACSS2, which converts acetate into acetyl-CoA, thereby increasing histone acetylation and promoting transcription of pro-survival genes (Fig. 4J). Similarly, uridine supplementation partially rescues cell death by supporting the pyrimidine salvage pathway. Uridine is directly converted to UMP by UCK1/2, restoring nucleotide balance and sustaining cell proliferation and tumor growth. Importantly, this uridine-mediated rescue occurs independently of TCA cycle intermediate restoration and does not involve alterations in cytosolic acetate metabolism. Collectively, these findings demonstrate that ACSS1 sustains tumor growth by supporting mitochondrial TCA cycle intermediates and pyrimidine precursor production, while also revealing metabolic vulnerabilities that may be therapeutically exploited upon ACSS1 loss.

**Fig. 4.**
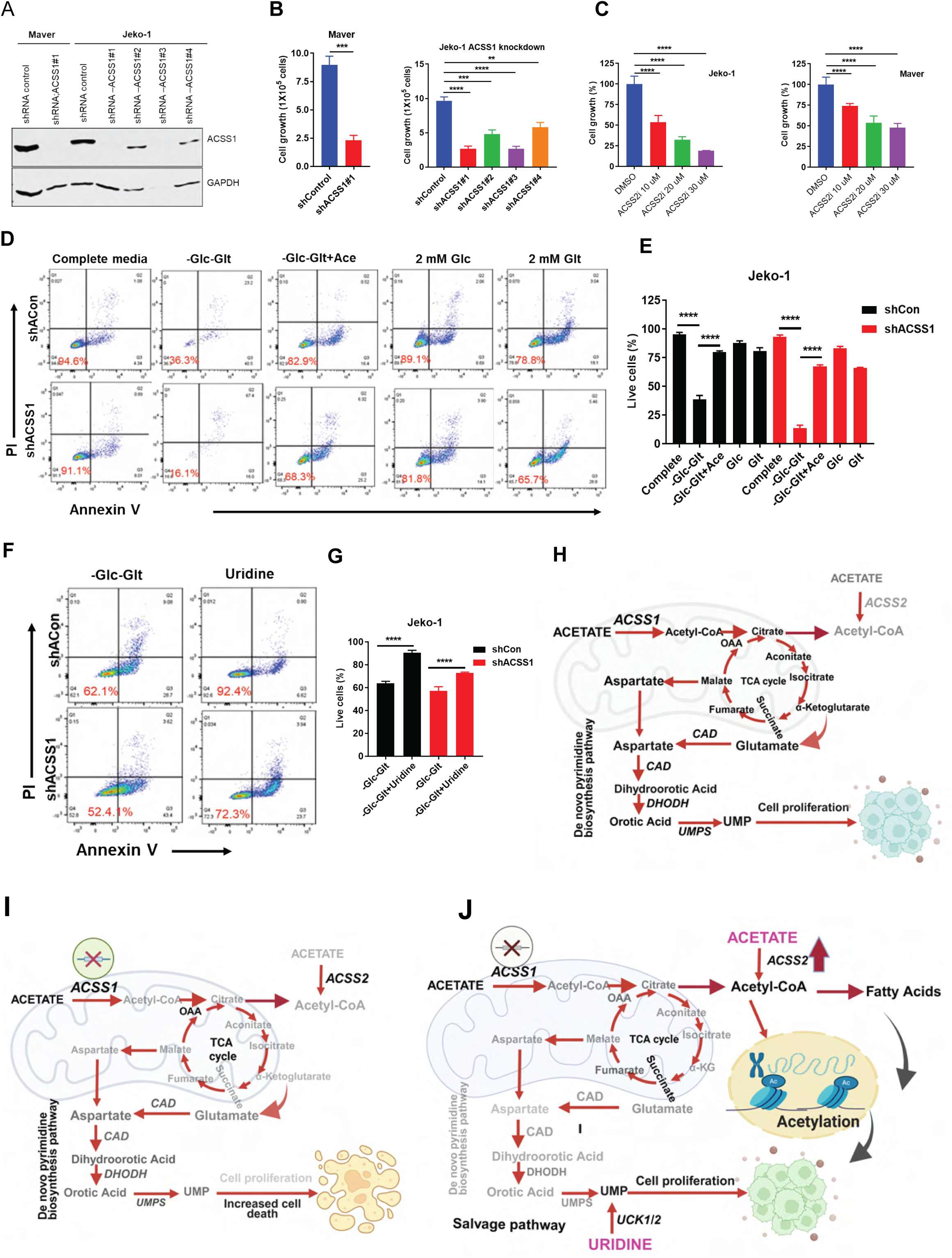
ACSS1 supports growth under nutrient restriction. **A**, Western blot analysis confirming efficient knockdown of ACSS1 in Maver and Jeko-1 cells using four independent shRNA clones; two clones exhibited near-complete protein depletion. **B**, ACSS1 knockdown (KD) significantly reduced cell growth, assessed by automated cell counting and trypan blue exclusion over 72 hours (P < 0.0001). **C**, Dose-dependent inhibition of cell growth by ACSS2 inhibitor (10–30 μM) in Jeko-1 and Maver cells cultured under standard conditions. **D, E**, Cell viability under metabolic stress (glucose/glutamine-free media + 1% dialyzed FBS) was markedly reduced in shACSS1 cells compared to shCon. Supplementation with 100 μM acetate significantly rescued viability in both groups, with a more pronounced effect in ACSS1 KD cells. **F, G**, Supplementation with 100 μM uridine rescued viability in both control and ACSS1 KD cells under nutrient-deprived conditions (P < 0.0001), implicating impaired pyrimidine biosynthesis in reduced cell survival. **H.** Schematic model illustrating ACSS1-mediated conversion of acetate to mitochondrial acetyl-CoA, fueling the TCA cycle and promoting the production of aspartate and glutamate to support de novo pyrimidine biosynthesis. The model highlights the metabolic advantage conferred by ACSS1 overexpression in sustaining nucleotide synthesis and tumor growth. **I.** Schematic model depicting impaired conversion of acetate to mitochondrial acetyl-CoA in ACSS1 knockdown cells. Reduced mitochondrial acetyl-CoA availability attenuates TCA cycle activity, leading to diminished production of aspartate and glutamate and consequent suppression of pyrimidine biosynthesis. **J.** Schematic model showing partial rescue of ACSS1-deficient cells. Acetate supplementation enhances compensatory ACSS2 activity in the cytosol, increasing acetyl-CoA production and histone acetylation to promote transcription of pro-survival genes. Uridine supplementation independently restores nucleotide balance by directly supporting UMP synthesis through the pyrimidine salvage pathway, thereby sustaining nucleotide production under ACSS1-depleted conditions.

**Fig. 5.**
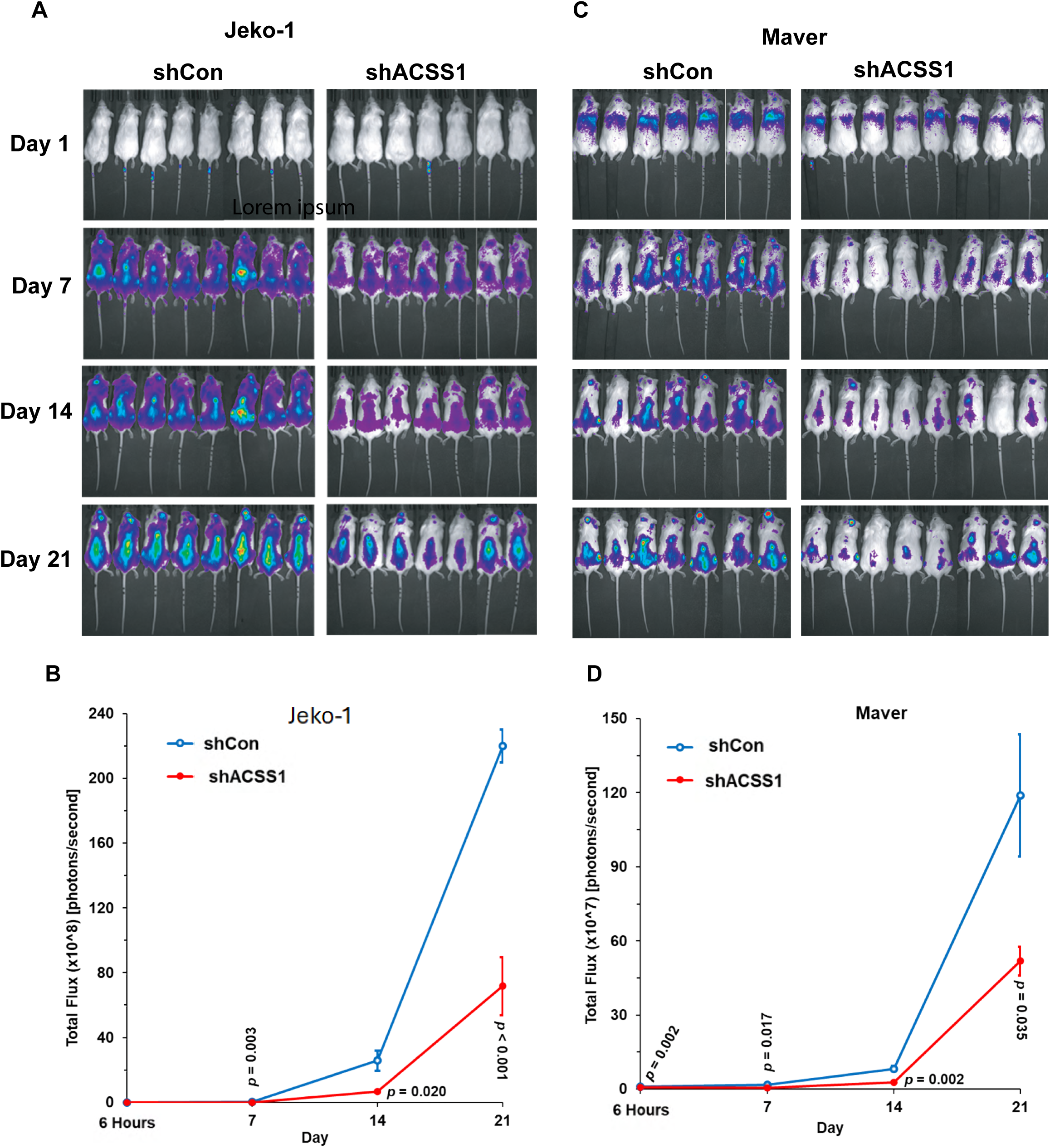
ACSS1 knockdown reduces tumor burden *in-vivo* in mantle cell lymphoma xenograft models. **A**. Representative bioluminescence imaging (BLI) of NSG mice injected with luciferase-expressing JeKo-1 cells transduced with control shRNA (shCon) or ACSS1-targeting shRNA (shACSS1). Tumor growth was monitored longitudinally by BLI. **B**. Quantification of tumor burden in the JeKo-1 model expressed as total photon flux (photons/sec) within defined regions of interest. Mice injected with shACSS1 cells exhibited significantly reduced tumor burden compared with shCon controls at days 14 and 21 post-inoculation (Welch’s two-sample t-test, two-tailed, p < 0.05). **C**. Representative BLI images of NSG mice injected with luciferase-expressing MAVER cells expressing shCon or shACSS1. **D**. Quantification of tumor burden in the MAVER model showing significantly reduced photon flux in the shACSS1 group at days 14 and 21 post-inoculation. Statistical significance was maintained after adjustment for multiple comparisons using the Holm–Bonferroni method.

### ACSS1 Knockdown Reduces *In Vivo* Tumor Burden in Mantle Cell Lymphoma Models

To evaluate whether ACSS1 promotes in vivo cancer growth during mantle cell lymphoma progression, we used Luciferase-expressing JeKo-1 and Maver cells expressing control shRNA (shCon) or shRNA targeting ACSS1 (shACSS1). These cells were injected into NSG mice, and tumor growth was monitored by bioluminescence imaging (BLI). The BLI imaging of Jeko-1 shCon and shACSS1 injected mice is shown (Fig.5A). Tumor burden was quantified as total photon flux (photons/sec) within defined regions of interest (Fig.5B). In the JeKo-1 model, mice injected with shACSS1 cells demonstrated a significant reduction in tumor burden compared with shCon controls at days 14 and 21 post-inoculation. BLI measurements revealed a progressive divergence between groups, with the shACSS1 cohort consistently exhibiting lower photon flux at these time points (Welch’s two-sample t-test, two-tailed p < 0.05).

Similarly, in the Maverl, ACSS1 knockdown resulted in significantly reduced tumor burden at days 14 and 21 compared with shCon-injected mice. The reduction in photon flux in the shACSS1 group was consistent across animals and remained statistically significant following adjustment for multiple comparisons using the Holm–Bonferroni method (Fig. 5C, D). These findings demonstrate that ACSS1 suppression significantly impairs in vivo tumor progression in both JeKo-1 and MAVER mantle cell lymphoma models.

## Discussion

In this study, we identify ACSS1 as a key metabolic enzyme that enables mantle cell lymphoma (MCL) cells to utilize acetate as a carbon source under nutrient-limited conditions. By integrating TCGA analysis, ^13^C-acetate isotope tracing metabolomics, and functional assays, we show that ACSS1 is highly expressed in MCL and other hematological malignancies and contributes to metabolic flexibility by supporting mitochondrial acetate metabolism, particularly when glucose and glutamine are scarce.

Our data show that MCL cell lines with high ACSS1 expression, such as Jeko-1 and Maver, demonstrate increased incorporation of ^13^C-acetate into TCA cycle intermediates, pyrimidine precursors, and downstream nucleotides, compared to RL cells with low ACSS1 expression. This suggests a model in which ACSS1 acts as a mitochondrial gatekeeper for acetate utilization, enabling continued energy production and biosynthesis in adverse nutrient environments. These findings are in agreement with prior studies in AML and melanoma models, where ACSS1 overexpression was linked to increased mitochondrial incorporation of acetate-derived carbon into the TCA cycle(23).

Our immunohistochemical analysis of patient-derived tumor samples confirms that ACSS1 is frequently expressed in MCL and DLBCL tissues, consistent with publicly available RNA-seq datasets from hematologic and solid tumors. Notably, we observed heterogeneous ACSS1 expression within and across lymphoma subtypes, suggesting metabolic diversity and differential reliance on acetate metabolism in the tumor microenvironment. ACSS1 expression correlated with increased incorporation of ^13^C-labeled acetate into TCA cycle intermediates and pyrimidine pathway metabolites, highlighting its role in maintaining biosynthetic capacity under conditions of glucose and glutamine deprivation.

Importantly, inhibiting ACSS1 impairs this metabolic flexibility. Our results show that knocking down ACSS1 markedly decreases ^13^C-acetate integration into TCA cycle intermediates, dihydroorotate, and orotate, which results in reduced pyrimidine production and diminished cell viability when nutrients are limited. These effects were partly restored by adding exogenous acetate or uridine, confirming that ACSS1 supports nucleotide synthesis through mitochondrial acetate metabolism. Additionally, oxygen consumption rates (OCR) depend on ACSS1 expression, showing that acetate oxidation via ACSS1 contributes to mitochondrial respiration and energy generation. Overall, our results identify ACSS1 as a key regulator of mitochondrial acetate metabolism in MCL. Unlike the better-understood cytosolic enzyme ACSS2, which promotes lipid biosynthesis and histone acetylation(24–26). ACSS1 specifically directs acetate into mitochondrial oxidative pathways and biosynthesis(14). In an earlier report, the role of Mitochondrial ACSS1-K635 acetylation knock-in mice shows altered liver lipid metabolism on a ketogenic diet(27). In another report, ACSS1-K635 acetylation knock-in mice exhibit altered metabolism, cellular senescence, and nonalcoholic fatty liver disease(3). This highlights the significance of compartmentalized acetate metabolism in helping cancer cells survive under metabolic stress.

Furthermore, our in vivo bioluminescence imaging confirmed that ACSS1-mediated acetate metabolism regulates tumor burden in mantle cell lymphoma.

Metabolic reprogramming is a hallmark of mantle cell lymphoma (MCL), yet the role of mitochondrial acetate metabolism in disease progression remains poorly defined. In this study, we demonstrate that suppressing ACSS1 significantly impairs in vivo tumor growth in two independent MCL xenograft models. Using luciferase-expressing JeKo-1 and MAVER cells, we show that ACSS1 knockdown results in a marked and sustained reduction in tumor burden at days 14 and 21 post-engraftment, as measured by longitudinal bioluminescence imaging (Fig. 5). The consistent attenuation of tumor growth across both models supports a functional role for ACSS1 in promoting MCL progression in vivo. Reduced tumor growth following ACSS1 suppression likely reflects impaired mitochondrial metabolism, limiting nucleotide biosynthesis and proliferation under in vivo conditions. The delayed divergence in tumor burden suggests that ACSS1 is particularly important as metabolic demand increases during tumor expansion.

These findings position ACSS1 as a potential metabolic vulnerability in MCL, distinct from cytosolic ACSS2, and suggest that targeting mitochondrial acetate metabolism could impair tumor growth. Future studies should clarify whether the effects of ACSS1 loss are mediated primarily through energy production, nucleotide synthesis, or other biosynthetic pathways, and assess potential compensatory mechanisms. Evaluating ACSS1 expression in primary MCL samples and its association with disease severity is also important to establish its translational relevance. In summary, ACSS1 promotes MCL progression in vivo, identifying mitochondrial acetate metabolism as a promising therapeutic target.

Therapeutically, these findings open new avenues for targeting metabolic dependencies in MCL. Inhibiting acetate metabolism, either by directly inhibiting ACSS1 or by limiting acetate availability, may hinder tumor cell survival, particularly in nutrient-poor microenvironments such as the tumor core or during treatment-induced metabolic stress. Additionally, ACSS1 expression could serve as a biomarker to stratify tumors by their dependence on mitochondrial acetate metabolism. Our results also suggest therapeutic implications. Tumors with high ACSS1 expression may be selectively vulnerable to strategies that disrupt acetate metabolism or target downstream biosynthetic pathways, such as pyrimidine synthesis.

In summary, we provide evidence that ACSS1-driven acetate utilization supports pyrimidine biosynthesis and cell survival under metabolic stress in MCL cells. These findings advance our understanding of metabolic plasticity in lymphoid malignancies and highlight ACSS1 as a potential metabolic vulnerability with therapeutic relevance.

## Materials and Methods

ACSS1 (Cat#17138-1-AP,RRID:AB_2289182), ACLY (Cat# 67166-1-Ig; RRID:AB_2882462), GAPDH (Cat# 60004-1-Ig;RRID:AB_2107436), DHODH ( Cat# 14877-1-AP, RRID:AB_2091723), CAD (Cat# 16617-1-AP, RRID:AB_2878289) and VDAC1 (Cat# 66345-1-Ig, RRID:AB_2881725) from Proteintech Inc., ACSS2 (Cat#3658S; RRID:AB_2222710) from Cell Signaling Technology. ^13^C2-Sodium acetate (Cat#CLM-440-1) from Cambridge Isotope Laboratories. The special media, D-Glucose-free RPMI 1640 (REF 11879-020), was obtained from Gibco. RPMI 1640 (-) Glutamine free and (-) D-Glucose free media were obtained from BI, Biological Industries, Israel Beit-Haemek. Uridine (Sigma, catalog#U3750-1G).

### MCL and DLBCL cell lines

Cell lines from different lymphomas were used, including MCL-RL(28), a cell line derived from a patient with MCL at the University of Pennsylvania, Philadelphia, PA, JeKo-1, Maver, Rec-1, and Granta519 for MCL; OCI-LY1, OCI-LY4, OCI-LY8, and TOLEDO for DLBCL; All commercially available cell lines were obtained from the American Type Culture Collection (ATCC). All the cell lines were regularly tested for Mycoplasma contamination using Mycoplasma detection kits from Thermo Fisher Scientific and authenticated. All cells were grown in RPMI medium supplemented with 10% FBS and 1% penicillin/streptomycin under a humidified 37°C/5% CO_2_ incubator. The cell lines are validated for authenticity and routinely tested for mycoplasma contamination.

### Nutrient starvation in cell culture

For nutrient-deprivation experiments, cells were washed once with phosphate-buffered saline. The glutamine-free medium was made with RPMI devoid of glutamine and glucose and retained the same nutrient concentrations as standard RPMI with dialyzed 10% FBS and 25 mM Glucose. A glucose-free medium was made with RPMI devoid of glutamine and glucose and retained the same nutrient concentrations as standard RPMI with dialyzed 10% FBS and 2 mM Glutamine. Complete RPMI was prepared by supplementing with 2 mM L-Glutamine and 25 mM Glucose to RPMI devoid of glutamine and glucose. For acetate supplementation, Sodium acetate at a final concentration of 5 mM was supplemented to Glutamine-free or Glucose-free media or both Glutamine/Glucose-free media.

### ACSS1 and mitochondrial fractionation

For mitochondria isolation, 50×10^6^ cells of Jeko-1 and Maver cell lines were harvested, and mitochondria were isolated according to the instructions of the product manual (catalog # 89874, Thermofisher). The protein estimation was measured, and an equal amount of protein was separated on a 10% NuPAGE gel and processed for western blotting as described below.

### ACSS1 knockdown (KD) in MCL cell lines

Jeko-1 and Maver cell lines were transduced with the scramble PLKO.1-PURO NON-TARGET CONTROL (SHC016V-1EA) and ACSS1 shRNA, TRCN0000436733, TRCN0000045380, TRCN0000045381, TRCN0000424197 and TRCN0000423168 all purchased from Millepore-Sigma-Aldrich. The ready-to-transduce lentivirus particles were directly mixed with 100,000 cells (Jeko-1 and Maver) in the presence of polybrene (catalog#TR-1003-G, Millepore Sigma) in a 96-well plate. The media was replaced with fresh RPMI after 16 hours of incubation. The cells were cultured for an additional 3 days and then selected with puromycin.

### In Vivo Bioluminescence Imaging

For in vivo bioluminescence imaging (BLI) studies, JeKo-1 and MAVER mantle cell lymphoma cell lines stably expressing either control shRNA (shCon) or shRNA targeting ACSS1 (shACSS1) were transduced in vitro with the Click Beetle Green luciferase reporter prior to inoculation to enable longitudinal tumor monitoring. For the JeKo-1 model, 0.7 × 10⁶ cells (shCon or shACSS1) were injected per mouse. For the MAVER model, 1.6 × 10⁶ shCon cells or 1.8 × 10⁶ shACSS1 cells were injected per mouse. Cells were administered via lateral tail vein injection into NSG mice. Tumor burden was quantified by measuring total photon flux (photons/sec) within predefined regions of interest (ROIs). Bioluminescence signals were collected longitudinally and used to assess tumor progression for each animal. Group means and standard errors of the mean (SEM) were calculated from individual animal measurements using standard propagation-of-error formulas. Statistical comparisons between groups were performed using Welch’s two-sample t-test applied to per-animal values. All tests were two-tailed. When multiple comparisons were conducted, p-values were adjusted using the Holm–Bonferroni method.

### Western blotting

Cells were harvested and processed for western blotting as previously described with respective antibodies; blots were developed using the Ibright 1500 imaging system (Thermo Scientific)(29).

### Reverse transcription quantitative polymerase chain reaction

MCL-RL, Jeko-1, and Maver, along with Jeko-1 shCon or shACSS1 cells, were seeded in six-well plates up to 48 hours. RNA isolation and DNase treatment were performed according to the kit protocol (Qiagen #74104, #79254). The RNA concentration was measured, and 1 μg was used for cDNA synthesis, which was performed with SuperScript IV (Invitrogen #18091050). For the quantitative reverse transcription polymerase chain reaction (qRT-PCR), 1 μL of cDNA was utilized. Each reaction contained: 1 μL of cDNA, 5 μL of SYBR Green mix (Applied Biosystem, A25742), 0.5 μL of forward primer (from a 10 mM stock), 0.5 μL of reverse primer (from a 10 mM stock), and 3 μL of dH2O. Each condition was tested in technical triplicate. Primers used are as follows,

### Primer Sequence 5’ to 3’

ACSS1, Forward- TTG AGA GCA CCC CAG TTT ATC; Reverse- GCA TCA CCG TAT TTC AGC AAC ACSS2, Forward- GGA TCA CTG GTC ATT CCT AC; Reverse- GTG CTG TGT AGA ACT TGG TC CAD, Forward- AGT GGT GTT TCA AAC CGG CAT; Reverse- CAG AGG ATA GGT GAG CAC TAA GA and DHODH, Forward-CCACGGGAGATGAGCGTTTC; Reverse-CAGGGAGGTGAAGCGAACA

### Immunohistochemistry (IHC) of MCL, DLBCL, and CLL patient-derived biopsy samples

We followed the protocols approved by the Institutional Review Board (IRB) of Fox Chase Cancer Center. Frozen DLBCL and MCL patient tissue samples were isolated from IRB-exempt discarded specimens of patients with diffuse large B-cell lymphoma (DLBCL) obtained through the Fox Chase Cancer Center Bio Specimen Repository. Human tissue microarrays for patients with MCL (*n* = 22), DLBCL (*n* = 28) and CLL (n=7) were used to examine ACSS1. The expression of ACSS1 was analyzed according to the pathology grade of the samples. Tissues were collected and fixed in 10% phosphate-buffered formaldehyde (formalin) for 24-48 hrs, dehydrated and embedded in paraffin. Hematoxylin and eosin (H&E) stained sections were used for morphological evaluation and unstained sections for IHC studies. IHC staining was performed on a VENTANA Discovery XT automated staining instrument (Ventana Medical Systems) using VENTANA reagents according to the manufacturer’s instructions. Slides were de-paraffinized using EZ Prep solution (cat # 950–102) for 16 min at 72 °C. Epitope retrieval was accomplished with CC1 solution (cat # 950–224) at high temperature (eg, 95–100 °C) for 32 min. Rabbit primary antibodies (ACSS1: 1:200, ACSS1 (Cat#17138-1-AP, RRID :AB_2289182) were tittered with a TBS antibody diluent into user-fillable dispensers for use on the automated stainer. Immune complex was detected using the Ventana OmniMap anti-rabbit detection kit (760–4311) and developed using the VENTANA ChromMap DAB detection kit (cat # 760-159) according to the manufacturer’s instructions. Slides were then counterstained with hematoxylin II (cat # 790-2208) for 8 min, followed by Bluing reagent (cat # 760-2037) for 4 min. The slides were dehydrated with ethanol series, cleared in xylene, and mounted. As a negative control, the primary antibody was replaced with normal rabbit IgG to confirm the absence of specific staining. Stained slides were scanned using an Aperio ScanScope CS 5 slide scanner (Aperio, Vista, CA, USA). Scanned images were then viewed and captured with Aperio’s image viewer software (ImageScope, version 11.1.2.760, Aperio).

### Cell proliferation and cell death analysis

For measuring cell proliferation, MCL Jeko-1 and Maver parental cell lines were seeded at a density of 2 × 10^4 cells per well in 96-well plates (Corning, NY, USA). Afterwards, the cells were treated with different concentrations of the ACSS2 inhibitor and incubated for 72 hours. After incubation, the cells were counted using the trypan blue method with a hemocytometer.

### Cell Culture Conditions under Nutrient Deprivation

Jeko-1 and Maver lymphoma cell lines shCon and shACSS1 were cultured under various nutrient conditions to assess the role of ACSS1 in supporting cell viability during metabolic stress.

Cells were seeded at a density of 2 × 10⁵ cells/mL and cultured for 72 hours under the following conditions: Complete RPMI medium supplemented with 10% FBS, Glucose- and glutamine-free RPMI supplemented with 1% dialyzed FBS, Glucose- and glutamine-free RPMI supplemented with 1% dialyzed FBS and 1 mM sodium acetate, Glucose- and glutamine-free RPMI supplemented with 1% dialyzed FBS and 2 mM glucose, and Glucose- and glutamine-free RPMI supplemented with 1% dialyzed FBS and 2 mM glutamine For pyrimidine depletion and rescue experiments, shACSS1 and shCon cells were cultured in glucose- and glutamine-free RPMI supplemented with 1% dialyzed FBS, in the presence or absence of 100 μM uridine (Cat# U3750-1G, Sigma-Aldrich), a pyrimidine nucleoside metabolite.

Following incubation, cell viability was assessed using the protocol as described in the kit for Annexin V / Propidium Iodide (PI) Staining ((BD Biosciences Cat# 556547, RRID:AB_2869082BD) and flow cytometry for cell viability assay.

### Metabolomics analysis

Metabolomics analyses were conducted at The Wistar Institute Proteomics and Metabolomics Shared Resource with slight modifications as previously described(30). For ^13^C_2_-acetate tracing experiments, the cells were cultured in glucose- and glutamine-free RPMI-1640 media supplemented with 500 µM labeled ^13^C_2_-acetate (Cambridge Isotope Laboratories Inc.), 10 mM unlabeled Glucose for 16 hours in triplicate. The cells were counted using the trypan blue method, and an equal number of cells (10×10^6^ cells/condition) were spun down at 1,000 rpm for 5 minutes. The cell pellet was washed with cold PBS and snap-frozen in liquid nitrogen. Unlabeled controls were generated for each cell line condition.

### Oxygen consumption rate (OCR) measurement by Seahorse analysis

The MCL cell lines (Jeko-1 shControl (control), Jeko-1 shACSS1 knockdown (KD), Maver shControl (control), Maver shACSS1 knockdown (KD) were exposed to either -glucose (Glc) or 2 mM acetate (ACE) for 1 hour and then thoroughly examined for mitochondrial respiration using the Seahorse-based method. The XF Mitochondrial stress test kit protocols (Agilent) were followed using the Agilent Seahorse XFe96 analyzer per the manufacturer’s instructions. In brief, cells were seeded in Seahorse XF 96-well plates at a density of 1.2x 10^5^ cells per well with Assay Base media, supplemented with 10 mM Glucose, 1 mM Sodium Pyruvate, and 2 mM Glutamine. The cell cultures were allowed to equilibrate for 1 hour at 37 °C in a no-CO2 incubator. The Oxygen Consumption Rate was analyzed under basal conditions and after treatment with different drugs, including 1 µM oligomycin A, 2 µM FCCP, and 0.5 µM Rot/AA. After the analysis, the medium was removed, and the cells were suspended in RIPA lysis buffer. The protein content of the cell lysates was then measured using the Bradford assay and used to normalize the respiratory parameters. Samples were analyzed with at least 6 technical replicates. The data were assessed using XF Wave Software (Seahorse Bioscience, Agilent).

### Statistics and reproducibility

The student’s unpaired t-test was used to analyze differences in quantitative (q)-PCR expression analysis, and isotope-labeled metabolites compared with MCL-RL cell line vs. Jeko-1 and Maver in three ^13^C-isotope tracing experiments. For ACSS1 knockdown (KD) studies in Jeko-1 and Maver cell lines, an unpaired t-test was used to analyze differences between control vs. KD cells ^13^C-acetate isotope-labeled metabolites. The student’s t-test was used to analyze differences in metabolites. p-values equal to or less than 0.05 were considered statistically significant without being adjusted for multiple comparisons. The statistical analysis was performed using GraphPad Prism 9.0. software.

## Data availability

We confirm that all relevant data and methods are included in the main Article and the Supplementary Information section.

## Acknowledgments

We are grateful to Dr. Erica Golemis, Senior Associate Dean of Research at Lewis Katz School of Medicine, Temple University Health System, and Professor and Chair, Cancer and Cellular Biology, Fox Chase Cancer Center, Philadelphia, for critically reading the manuscript and providing feedback to improve the paper. We thank the IHC core facility members at Fox Chase Cancer Center for providing excellent IHC staining. We thank Dr. Andrey Efimov, Bio Imaging Facility, Fox Chase Cancer Center, for confocal imaging. We created the figures in this manuscript using the Biorender program..

## Funding

The funding for the startup comes from Fox Chase Cancer Center to MW. The Wistar Institute Proteomics and Metabolomics Shared Resource is supported by NIH Cancer Center Support Grant CA010815. The metabolomics analysis was performed on a Thermo Q-Exactive HF-X mass spectrometer purchased with NIH grant S10 OD023586.

## Author Contributions

Conceptualization: J.B. Writing original draft: J.B. Data curation: J.B. Methodology: J.B., A.R.G. and N.V.S. Formal Analysis: J.B., A.R.G., and D.R. Investigation: J.B., C.L., S.W., D.R., O.M., N.V.S., V.S.M., P.J., M.E., R.N., K.Q.C., D.S.N., P.K.G., and K.N. Visualization: J.B., A.R.G., and C.L. Validation: J.B., A.R., D.R. and N.V.S. Resources: H.B., and K.E.W. Supervision: J.B. Writing, review and editing: J.B., A.R.G, C.L., K.E.W. and M.A.W. Funding acquisition: M.A.W. Read and approved of the manuscript: J.B., A.R.G., C.L., S.W., D.R., O.M., N.V.S., V.S.M., P.J., M.E., K.C., P.L., R.N., H.G., K.E.W., and M.A.W.

## Competing interests

All authors declare that they have no competing interests.

## Data and materials availability

All data needed to evaluate the conclusions in the paper are present in the paper and/or the Supplementary Materials.

---

**Supplementary Fig. 1.**
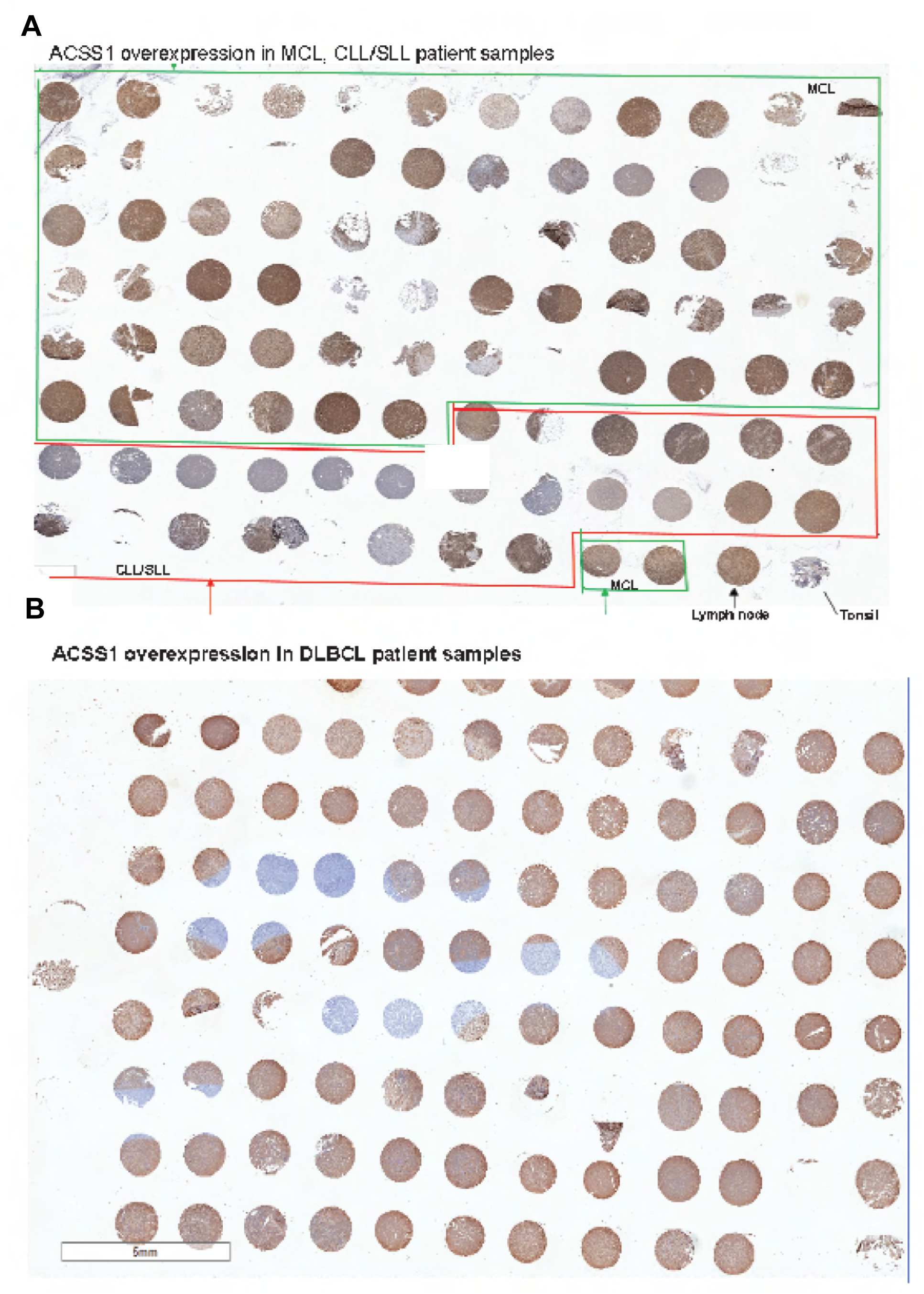
Overview of tissue microarrays (TMAs) and ACSS1 expression in lymphoma. **A**, Low-magnification overview of the mantle cell lymphoma (MCL) TMA block used for immunohistochemical analysis of ACSS1 expression. **B**, Representative IHC images showing ACSS1 staining in diffuse large B-cell lymphoma (DLBCL) patient samples (n = 13). ACSS1 expression was detected in 7 cases (53.8%).

**Supplementary Fig. 2.**
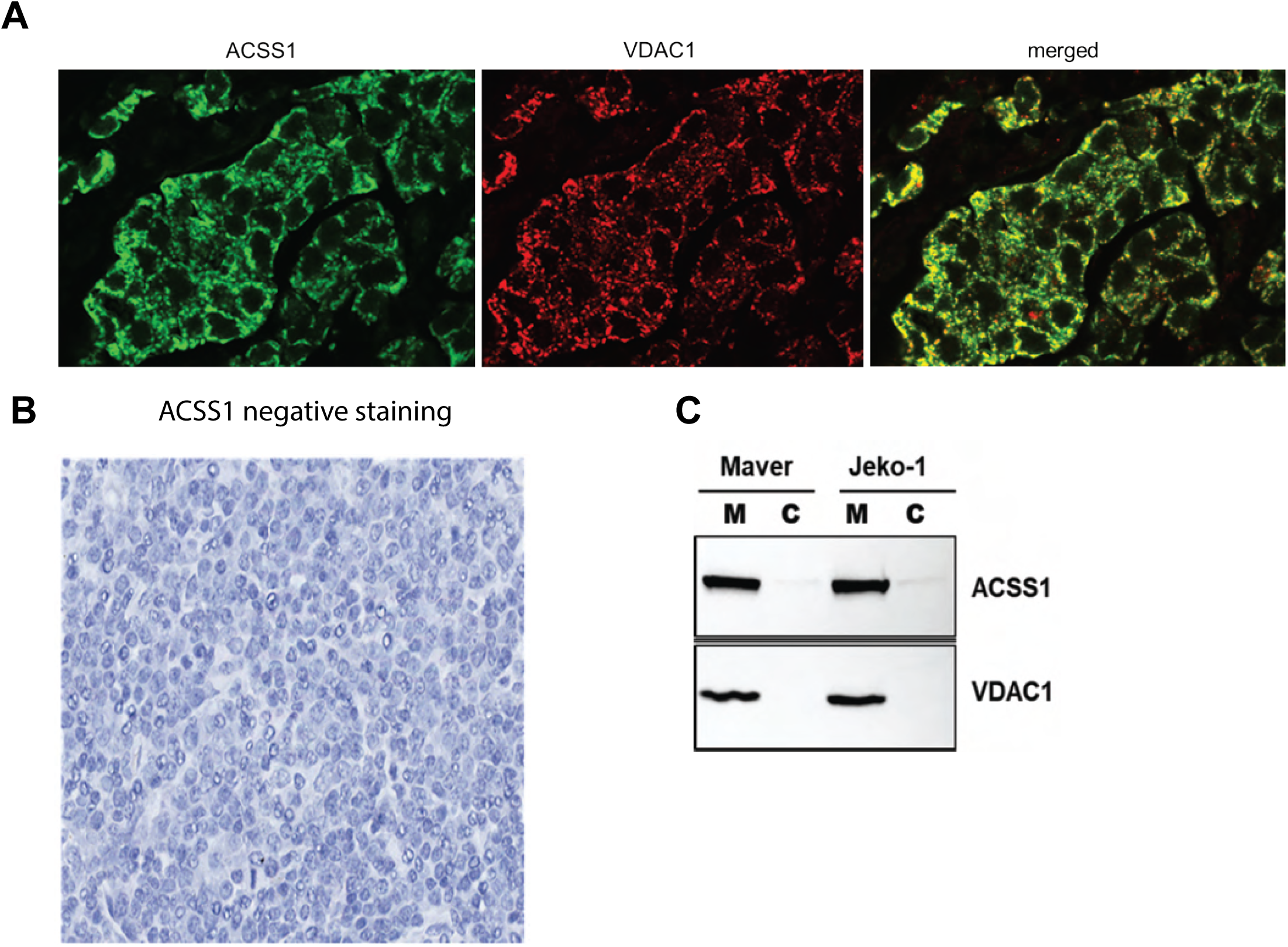
Validation of ACSS1 antibody specificity and mitochondrial localization. **A, B**, Immunofluorescence analysis of a breast cancer tissue section stained with antibodies against ACSS1 (green) and the mitochondrial marker VDAC1 (red), with nuclei counterstained using DAPI (blue). Confocal microscopy shows co-localization of ACSS1 and VDAC1 in positive control tissue. **C**, Negative control showing absence of ACSS1 signal in breast cancer tissue, confirming antibody specificity. **D**, Subcellular fractionation and immunoblot analysis of MCL cell lines (Maver and Jeko-1) demonstrate enrichment of ACSS1 in mitochondrial fractions. VDAC1 serves as a mitochondrial marker.

**Supplementary Fig. 3.**
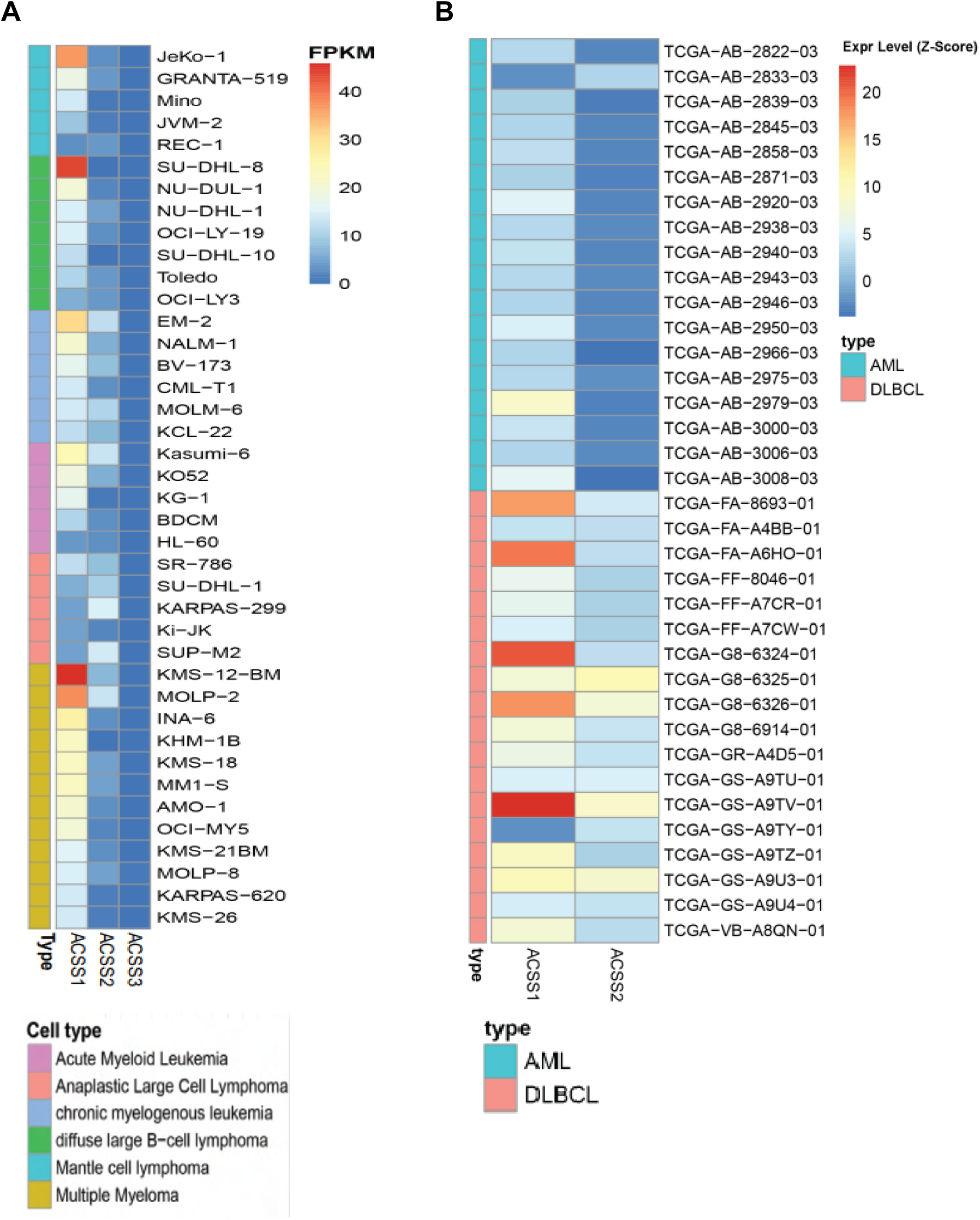
ACSS1 is selectively expressed in non-Hodgkin lymphoma models. **A**, RNA-seq–based expression analysis of acetyl-CoA synthetase family members (ACSS1, ACSS2, ACSS3) across hematological cancer cell lines using the Expression Atlas database. Data includes models of mantle cell lymphoma (MCL), diffuse large B-cell lymphoma (DLBCL), chronic lymphocytic leukemia (CLL), acute myeloid leukemia (AML), and anaplastic large cell lymphoma (ALCL). **B**, ACSS1 expression is elevated in non-Hodgkin lymphoma (NHL) cell lines and patient-derived samples, whereas ACSS3 expression is consistently low or undetectable (FPKM <1) across all profiles.

**Supplementary Fig. 4.**
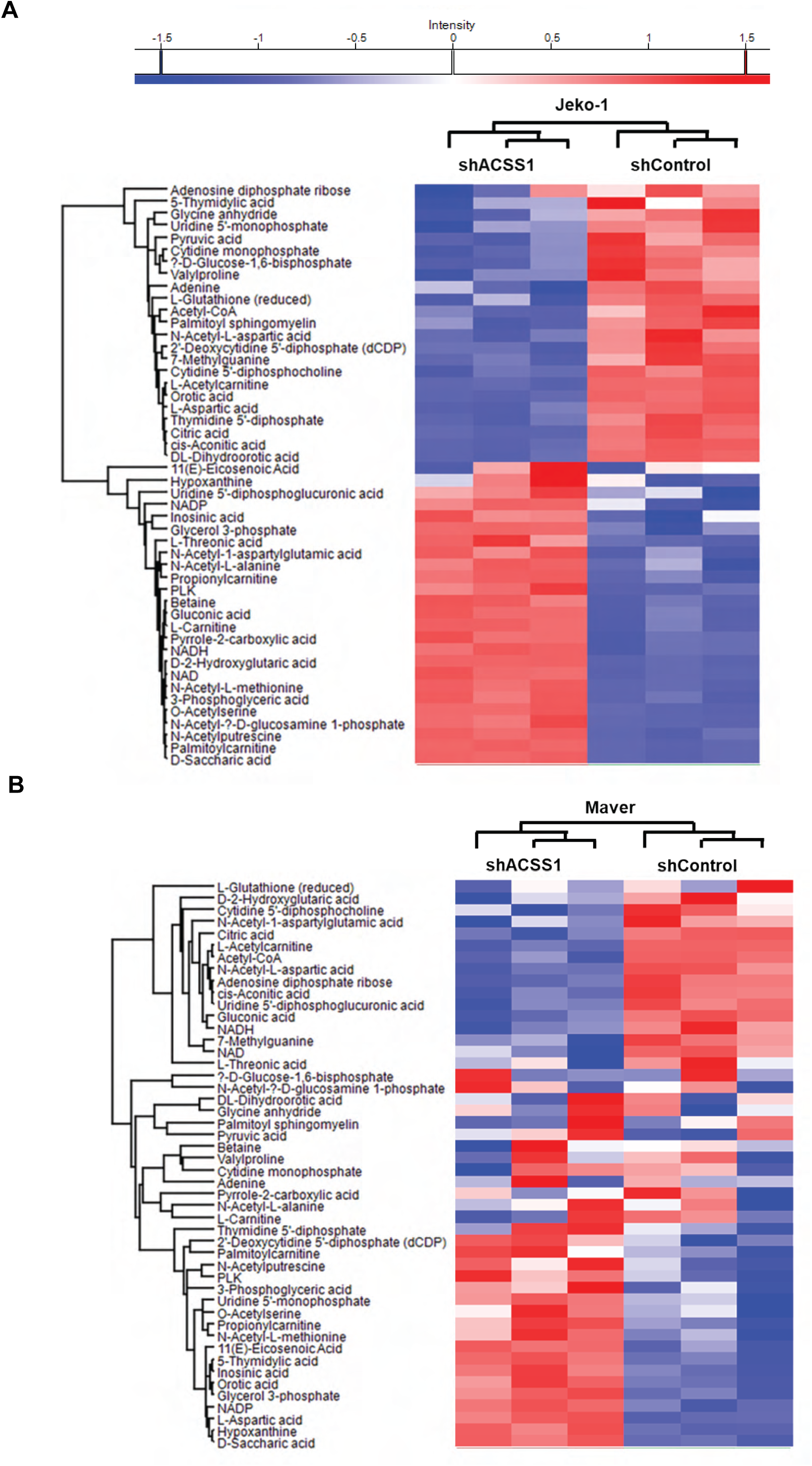
ACSS1 knockdown alters pyrimidine-associated metabolites under acetate-reliant conditions. **A, B,** Heatmap analysis of metabolomic profiling of shACSS1 and shControl Jeko-1 and Maver cells cultured in glucose- and glutamine-free media supplemented with 1% dialyzed FBS and ¹³C-acetate. Of the 206 metabolites analyzed, 46 showed significant changes (fold change >1.5, P < 0.05), with a majority enriched in the pyrimidine biosynthesis pathway. Data represents results from two independent cell lines (Jeko-1 and Maver), each with triplicate biological replicates.

## Notes

### Competing Interest Statement

The authors have declared no competing interest.

### Summary of Updates

Figure 5 is added. Importantly, in vivo studies using luciferase-labeled JeKo-1 and Maver mantle cell lymphoma xenografts demonstrate that ACSS1 knockdown significantly suppresses tumor growth. NSG mice injected with ACSS1-silenced cells exhibit a marked reduction in tumor burden, as measured by bioluminescence imaging and total photon flux, with significant differences observed at days 14 and 21 post-injection.

